# Simultaneous detection of protozoa, bacteria, and viruses from environmental water through membrane-adsorption followed by direct nucleic acid extraction

**DOI:** 10.1101/2025.08.07.669200

**Authors:** Shotaro Torii, Masaaki Kitajima, Eiji Haramoto, Ryota Gomi, Kumiko Oguma, Hiroyuki Katayama

**Affiliations:** Department of Urban Engineering, School of Engineering, The University of Tokyo, 7-3-1 Hongo, Bunkyo-ku, Tokyo 113-8656, Japan; Research Center for Water Environment Technology, School of Engineering, The University of Tokyo, 2-11-16 Yayoi, Bunkyo-ku, Tokyo 113-0032, Japan; Interdisciplinary Center for River Basin Environment, University of Yamanashi, 4-3-11 Takeda,Kofu, Yamanashi, 400-8511, Japan

**Keywords:** Bacteria, Detection methods, EPISENS-M, Protozoa, Virus

## Abstract

Waterborne outbreaks caused by protozoan, bacterial, and viral pathogens continue to pose serious public health threats. However, comprehensive monitoring of all three major pathogen types remains uncommon, primarily due to the technical difficulty of simultaneously concentrating and characterizing these diverse microorganisms from a single water sample. Here, we evaluated a membrane adsorption followed by bead-beating-based direct nucleic acid extraction for its sensitivity in detecting waterborne pathogens. We further applied the optimized protocol to river water samples. Our results demonstrated that the use of magnesium chloride in combination with a mixed cellulose ester membrane filter (pore size: 0.8 μm) achieved the highest concentration efficiency for *Cryptosporidium parvum*, *Legionella pneumophila*, and murine norovirus (MNV), outperforming other methods utilizing the membrane with alternative pore sizes or membrane with another material (i.e., nylon). Furthermore, targeting rRNA of *C. parvum* instead of rDNA, or incorporating a preamplification step for the *L. pneumophila mip* gene and MNV, significantly improved the sensitivity. Notably, the 50% limit of detection for *C. parvum* using our method was comparable to that of traditional immunofluorescent antibody testing. Application of this method to river water further revealed a higher abundance of *Bacteroides* rRNA markers compared to their corresponding rDNA, underscoring the potential of rRNA-based targets for more sensitive bacterial detection. Importantly, when the target is genomic DNA or RNA, preamplification enables the use of a larger nucleic acids input volume, thereby enhancing sensitivity without introducing measurement bias. Overall, the combination of membrane adsorption and bead-beating-based direct nucleic acid extraction offers a sensitive, all-in-one approach for simultaneous detection of protozoa, bacteria, and viruses in environmental water. This method serves as an effective ‘screening’ tool for the presence of the three major types of waterborne microorganisms.

**Graphical Abstract:** 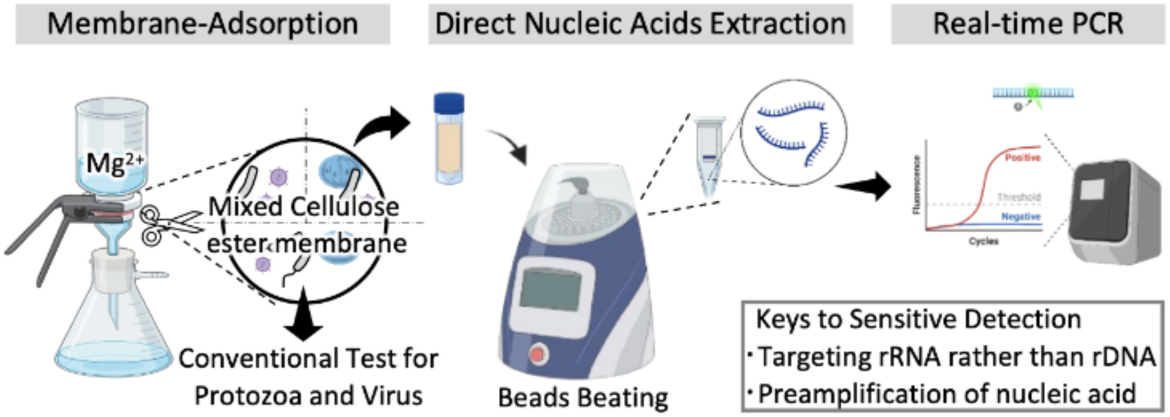

## 1. Introduction

Waterborne outbreaks of diseases caused by protozoa, bacteria, and viruses continue to pose significant public health risks (Gerdes et al., 2023). To assess microbial contamination in environmental and drinking water, fecal indicator bacteria (FIB), such as *Escherichia coli* and enterococci, are widely monitored as proxies for pathogenic organisms (Boehm et al., 2018). However, protozoa (e.g., *Cryptosporidium*) and viruses (e.g., norovirus) are not routinely monitored, despite their greater resistance to environmental stressors compared to bacteria (Boehm et al., 2019).

Protozoa, bacteria, and viruses differ markedly in size and biological characteristics. As a result, detection methods are typically developed separately for each group of microorganisms. This separation complicates efforts to simultaneously detect multiple pathogen types from a single water sample. Some studies have sought to address this issue by developing concentration methods capable of capturing all three microorganism groups, including ultrafiltration (Hill et al., 2007; Morales-Morales et al., 2003), skimmed milk flocculation (Gonzales-Gustavson et al., 2017), and membrane adsorption using negatively charged microfiltration membranes following addition of magnesium chloride (Haramoto et al., 2012). However, even with these approaches, subsequent detection procedures typically remained microorganism-specific.

Traditionally, protozoa are detected by immunofluorescent antibody tests (IFAT), while bacteria and viruses are characterized using culture-based methods. Although these approaches provide direct evidence of viable or infectious microorganisms, they are time-consuming, labor-intensive, and expensive. Furthermore, each test typically targets only a limited group of microorganisms, resulting in fragmented workflows that can burden water utilities. In contrast, molecular-based approaches using nucleic acid extraction followed by real-time PCR offer a more universal and scalable platform. Moreover, these methods can allow for more detailed microbial characterization. For example, real-time PCR products for *Giardia* spp. can be directly sequenced to assess whether the detected *Giardia* is human-infectious type (Jothikumar et al., 2021), whereas IFAT would require an additional, separate step for such differentiation.

Despite their versatility, molecular-based methods face two major limitations. First, efficient nucleic acid extraction is challenging, especially for protozoan (oo)cysts, which are notoriously difficult to lyse (Mthethwa et al., 2022; Nantavisai et al., 2007) without a labor-intensive pretreatments (e.g., several to fifteen cycles of freeze-thawing (Nichols and Smith, 2004; Wells et al., 2016)). Second, PCR assays typically use only a small fraction of the total nucleic acid extract (e.g., testing only several microliters out of tens of microliters), which reduces the effective sample volume (ESV) and overall sensitivity. As a result, limited studies have achieved simultaneous molecular-based detection of the three microorganisms from the same concentrate, with notable exceptions of (Gonzales-Gustavson et al., 2017; Hill et al., 2007). Moreover, it remains unclear, particularly for protozoa, whether the choice of the molecular-based methods achieves as high sensitivity as the traditional method (e.g., IFAT) and whether the additional methodological refinements could improve the sensitivity.

Recent advancements in molecular biology offer promising solutions to these challenges. Direct nucleic acid extraction via bead-beating has been shown to improve recovery efficiency, even for difficult samples such as *Cryptosporidium* (Hachimi et al., 2024), and the method is applicable to membrane filters through which water samples are passed (Ando et al., 2023; Nakaso et al., 2024). Moreover, introducing a preamplification step, a short conventional PCR performed before real-time PCR, can enhance sensitivity by increasing the amount of targeted template. This approach increases template abundance without compromising quantitative accuracy and enables the use of larger template nucleic acid volumes (e.g., up to 13.5 μL), thus increasing ESV (Ando et al., 2022). Therefore, leveraging these two technologies is promising for the simultaneous, highly sensitive detection of the three types of organisms.

Here, we optimized a membrane adsorption followed by bead-beating-based direct nucleic acid extraction methods and evaluated its sensitivity for detecting *Cryptosporidium parvum*, *Legionella pneumophila*, and murine norovirus (MNV) as model organisms in laboratory-prepared water samples. We compared detection sensitivities across different molecular processes and IFAT. As a proof of concept, the optimized method was applied to twelve river water samples to detect two protozoa (*Cryptosporidium* spp. and *Giardia* spp.), two bacteria (*L. pneumophila* and the HF183 16S ribosomal RNA (rRNA) gene cluster of members of *Bacteroides*), and norovirus genogroup II (NoV GII).

## 2. Materials and Methods

### 2.1. Optimization and sensitivity of the membrane-adsorption followed by direct nucleic acid extraction

#### 2.1.1. Microorganisms

Viable *C. parvum* at a concentration of 1.25 × 10^6^ oocysts/mL was purchased (Waterborne Inc., New Orleans, LA, USA) and stored at 4 ℃ prior to use. *L. pneumophila* was propagated at buffered yeast extract medium at 37 ℃ for 48 h as described elsewhere (Rattanakul and Oguma, 2018) and stored at −80 ℃. MNV S7-PP3 strain was propagated on RAW 264.7 cells, as described elsewhere (Kitajima et al., 2010). The genomic concentration of *L. pneumophila* and MNV stocks were quantified by (RT)-quantitative PCR (qPCR) applying gBlocks Gene Fragments (Integrated DNA technologies, Coralville, IA, USA), whose concentrations were verified using droplet digital PCR (ddPCR) (Bio-Rad Laboratories, Hercules, CA, USA) as described in Supporting Information (SI) and Table S1, as standards.

#### 2.1.2. Preparation of test water

For optimization of experimental conditions, a hundred milliliters of reverse osmosis (RO) water was spiked with 2.0 × 10^4^ oocysts of *C. parvum*, 2.0 × 10^5^ genome copies (gc) of *L. pneumophila*, and 7.2× 10^4^ gc of MNV. For the test of sensitivity, three types of test water were prepared by spiking different loads of *C. parvum*, *L. pneumophila*, and MNV into ten liters of distilled water. The loaded amount differs at three levels, ranging 0.5–50 oocysts for *C. parvum*, 5–500 gc for *L. pneumophila*, and 500–50000 gc for MNV. The lowest load for each organism was combined and added to one test water. The remaining two test waters were spiked with 10-fold and 100-fold higher loads, respectively.

#### 2.1.3. The concentration of the microorganisms by membrane-adsorption

For optimization of experimental conditions, the concentration was performed under two different conditions, including (i) the supplementation of the test water with magnesium chloride at a final concentration of 25 mM and (ii) the type of membrane filters used. We evaluated 47 mm mixed cellulose ester (MCE) membranes with different pore sizes (0.45–3.0 μm, each of which is referred to as 0.45-MCE, 0.8-MCE, 1.2-MCE, and 3.0-MCE, respectively; Merck Millipore, Billerica, MA, USA) and a 47 mm nylon net filter with a pore size of 0.45 μm, referred to as 0.45-Nylon (Merck Millipore).

For sensitivity evaluation, test water was supplemented with magnesium chloride at a final concentration of 25 mM and was filtered through a 90 mm 0.8-MCE. The membrane was then cut in half, and one half was used for nucleic acid extraction.

#### 2.1.4. Nucleic acid extraction

DNA and RNA were co-extracted with a slightly modified version of the Efficient and Practical virus Identification System with ENhanced Sensitivity for Membrane (EPISENS-M) (Ando et al., 2023; Nakaso et al., 2024). The membrane filter was inserted into Precellys 7 mL tubes (Bertin Technologies, Montigny-le-Bretonneux, France), where beads from RNeasy PowerWater Kit (Qiagen, Hilden, Germany), 850 μL of preheated buffer PM1, 150 μL of TRIzol (Thermo Fisher Scientific, Waltham, MA, USA), and 10 μL of β-mercaptoethanol (FUJIFILM Wako Pure Chemical Corporation, Osaka, Japan) were included. The tubes were then subjected to bead-beating five times at 10,000 rpm for 20 sec with 15 sec intervals using the Precellys Evolution tissue homogenizer (Bertin Technologies). After centrifugation at 10,000 × *g* for 3 min, 450 μL of the supernatant was transferred to a rotor adapter. Co-extraction of DNA and RNA was carried out with QIAcube Connect platform (Qiagen) using the automated program for RNeasy PowerMicrobiome Kit (Qiagen), where the inhibitor removal steps were included, but DNase treatment was omitted (Nakaso et al., 2024). DNA and RNA were eluted with 50 μL of DNase/RNase free water.

#### 2.1.5. Reverse transcription

A 7.5 μL of the extract was subjected to reverse transcription (RT) using the High-Capacity cDNA Reverse Transcription Kit (Thermo Fisher Scientific,) to obtain 15 μL of (c)DNA, following the manufacturer’s protocol.

#### 2.1.6. RT-preamplification

One-step RT-preamplification was performed to prepare a template for real-time PCR using iScript^TM^ Explore One-Step RT and PreAmp Kit (Bio-Rad) according to a previous study (Ando et al., 2022). Briefly, 13.5 μL of the nucleic acid extract was mixed with 16.5 μL of the RT-Preamp reaction mixture, containing 15.0 μL of SsoAdvanced Preamp Supermix, 0.6 μL of iScript Advanced Reverse Transcriptase, 0.6 μL of iScript Explore Reaction Booster, and 0.3 μL of primer mix containing 3 pmol each of forward and reverse primers of *mip* gene of *L. pneumophila* and MNV. Primers for *Cryptosporidium* spp. 18S rRNA gene were not included because our prior experiment suggests that the inclusion of the primers led to unspecific amplification from no template control (NTC) using nuclease-free water (data not shown). Nuclease-free water was used as a negative control for pre-amplification. Tenfold serial dilution of gBlocks Gene Fragments (5×10^4^ to 5×10^-1^ gc/μL), containing the amplified region of *mip* gene and MNV was also subjected to preamplification to generate a standard curve. The thermal cycle condition was 25 ℃ for 5 min, 45 ℃ for 60 min, and 95 ℃ for 3 min, followed by 10 cycles of 95 ℃ for 15 sec and 58 ℃ for 4 min, according to the manufacturer’s instructions.

#### 2.1.7. qPCR

qPCR was performed in a total reaction volume of 25 μL, which consisted of 2.5 μL of the template and 22.5 μL of the reaction mixture. The reaction mixture contained 12.5 μL of QuantiTect Probe PCR Master Mix (Qiagen), 10 pmol each of forward and reverse primer, and 5 pmol of probe. The primers and probes used in this study are according to previous studies (Jothikumar et al., 2008; Kitajima et al., 2010; Nazarian et al., 2008) and are listed in Table S2. PCR amplification was performed under the following thermal conditions: 50 ℃ for 2 min and initial denaturation at 95 ℃ for 15 min, followed by 45 cycles of denaturation at 95 ℃ for 15 sec and annealing and extension at 60 ℃ for 1 min. For NTC, nuclease-free water instead of template was included. Amplification data were analyzed by QuantStudio^TM^ Design & Analysis Software v1.5.2 (Thermo Fisher Scientific). The threshold of relative fluorescence intensity was adjusted to be 0.01 according to the recommendation of the mastermix manufacturer. If preamplified templates are used, preamplified gBlocks were used to generate a standard curve. Otherwise, ten-fold serial dilutions of gBlocks Gene Fragments (5×10^4^ to 5×10^-1^ gc/μL) were prepared and used to generate a standard curve. All the amplification that occurred above Ct value of 40 was regarded as non-detection.

### 2.2. Simultaneous detection of protozoa, bacteria, and viruses from surface water

#### 2.2.1. Sampling site, sample collection, and concentration

The downstream bank of the Tama River (N° 35.611540, E° 139.622275) was selected as a sampling site in this study, as an example of surface water heavily affected by microbial pollution. The Tama River flows through Tokyo and Kanagawa Prefectures. Its upstream section serves as a regional water supply, while its downstream section, including the sampling site, is heavily impacted by effluents from wastewater treatment plants and has not been used as a water source since 1970. Two-liter river water samples were collected monthly between May 2023 to April 2024, resulting in a total of 12 samples. Within one day of collection, one liter from each sample was concentrated as described in section 2.1.3. The filtered membranes were cut into quarters and stored at −20℃.

#### 2.2.2. Molecular quantification of protozoa, bacteria and viruses

A quarter piece of membrane filter was extracted as per section 2.1.4. A 7.5 µL of nucleic acid extract was subjected to RT as per section 2.1.5. A 13.5 µL of the extract was subjected to one-step RT-preamplification with forward and reverse primers of *mip* gene and NoV GII (Bernhard and Field, 2000; Green et al., 2014; Kageyama et al., 2003; Nazarian et al., 2008). Tenfold serial dilutions of gBlocks Gene Fragments (5×10^4^ to 5×10^-1^ gc/μL, verified by ddPCR), containing amplified region of *mip* gene and NoV GII, were also prepared to generate a standard curve. Real-time PCR was run according to section 2.1.7. For the assays targeting rRNA genes, including *Cryptosporidium* spp., *Giardia* spp. (Jothikumar et al., 2021), and HF183, both RT and non-RT samples were tested. For the *mip* gene of *L. pneumophila*, samples with RT, without RT, and with RT-preamplification were evaluated. In the case of NoV GII, which has genomic RNA, only samples processed with RT and with RT-preamplification were analyzed. The real-time PCR products of *Giardia* spp. were purified and submitted for nanopore sequencing using the Premium PCR sequencing service (Plasmidsaurus, Arcadia, CA, USA) as detailed in the SI.

#### 2.2.3. Enumeration of (oo)cysts of *Cryptosporidium* spp. and *Giardia* spp

Recovery of protozoa from mixed cellulose ester membranes was performed as described elsewhere (Haramoto et al., 2012), which reported a recovery efficiency of 49% for *Cryptosporidium* oocysts and 48.5% for *Giardia* cysts from river water. To elute protozoa attached to the membrane filters, another quarter was vigorously vortexed with 15 mL of an elution buffer (0.2 g/L of Na_4_P_2_O_7_·10 H_2_O, 0.3 g/L of EDTA trisodium salt trihydrate, and 0.1 mL/L of Tween 80) in a 50-mL centrifuge tube containing a stirrer bar. The eluate was then centrifuged at 1,100 × *g* for 10 min at room temperature, and the pellet was resuspended in 5 mL of phosphate-buffered saline (PBS)

The resuspensions were purified via immunomagnetic separation (IMS) using Dynabeads GC-Combo (Thermo Fisher Scientific) according to the manufacturer’s instructions. The immuno-separated samples were filtered through a hydrophilic PTFE membrane (pore size, 1 µm, Millipore).

Viable (oo)cysts were detected via an IFAT. Immunostaining was performed using EasyStain (Biopoint, Sydney, Australia) according to the manufacturer’s protocol. Briefly, the filter was stained with DAPI, followed by FITC-conjugated *Cryptosporidium* spp.- and *Giardia* spp.-specific antibodies. Enumeration was carried out using a Nikon Eclipse Ti2-E equipped with a Nikon DS10 (Nikon, Tokyo, Japan). The stained filter was visualized with the B excitation (wavelength of 450– 490 nm). Round-shaped fluorescent particles with a diameter of 4–6 μm were counted as *Cryptosporidium* spp. while oval-shaped fluorescent particles with a diameter of 5–8 μm and a width of 8–12 μm were counted as *Giardia* spp. Particles that fluoresced under G excitation (wavelength of 546 nm) were excluded as algae because chlorophyll in the algae fluoresces under this wavelength (Rodgers et al., 1995).

### 2.3. Data analysis

All data analyses were performed using R 4.3.0 (R Core Team, 2019). Analysis of variance (ANOVA) was conducted, followed by post hoc Tukey HSD tests using the {emmeans} package. Fisher’s exact test was performed with the function fisher.test. The load of microorganisms is fitted by the probit regression model via the glm function. The resulting model was then solved to determine the load corresponding to a 50% probability of detection (LOD_50_).

## 3. Results

### 3.1. Optimization of concentration

Initially, we optimized two experimental parameters: the membrane pore size and supplementation of the water sample with magnesium chloride at a final concentration of 25 mM. Note that the results obtained by qPCR (without RT) were used for assessing the recovery of *C. parvum* and *L. pneumophilia* in this section 3.1. For MNV, which has genomic RNA, the results obtained by RT-qPCR were used for recovery assessment.

The recovered microbial load was converted into recovery efficiency, and both metrics are presented in Figure 1. For *C. parvum*, the recovered load ranged from 5.3 to 5.7 log_10_ gc, corresponding to recovery efficiencies of 69% to 153%. In the case of *L. pneumophila mip* gene, the recovered load ranged from 4.7 to 5.1 log_10_ gc, with recovery efficiencies between 40% and 73%. Although these values did not differ significantly with or without magnesium chloride supplementation or among tested membranes (ANOVA, *P* > 0.05), the 0.8-MCE yielded the highest recoveries for *C. parvum* and *L. pneumophila*.

**Figure 1.**
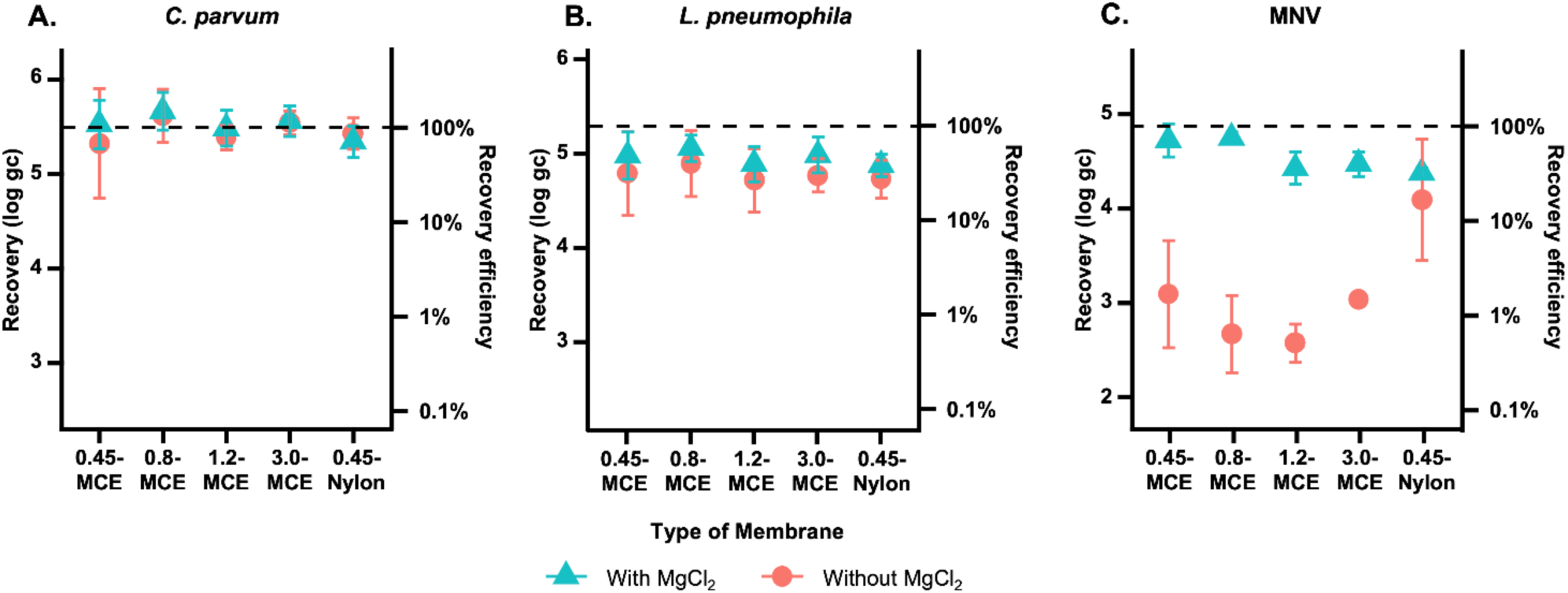
Recovery of *C. parvum* (A), *L. pneumophila* (B), and MNV (C) using various membrane filters and with or without magnesium chloride supplementation. The left axis indicates the recovered load (log_10_ gc), while the right axis shows the recovery efficiency relative to the inoculated load. Triangle symbols represent data obtained with magnesium chloride at 25 mM supplementation, while circular symbols represent data without magnesium chloride. Error bars indicate standard deviations (*n* = 3 for each experimental condition), and the dashed line represents the spiked microorganism load, corresponding to the recovery efficiency of 100%.

In contrast, the recovered load and recovery efficiency for MNV varied substantially, from 2.6 to 4.7 log gc and from 0.55% to 83%, respectively. Under magnesium chloride supplementation, the mean recovery efficiencies for MNV were 83% for the 0.8-MCE, 78% for the 0.45-MCE, 35% for the 0.45-Nylon, 39% for the 1.2-MCE, and 44% for the 3.0-MCE. Membrane type had a significant impact on recovery efficiencies, and the absence of magnesium chloride supplementation led to reduced recoveries across all membrane except for the nylon membrane (ANOVA, *P* < 0.05).

These results suggest that using magnesium chloride and a 0.8-MCE yields the highest concentration efficiency for *C. parvum*, *L. pneumophila*, and MNV. This optimized condition was used in all subsequent experiments.

### 3.2. Sensitivity

To determine the sensitivity of the optimized concentration method (i.e., using 0.8-MCE membrane with magnesium chloride supplementation), varying amounts of microorganisms were spiked into RO water and tested for detectability by real-time PCR. Table 1 summarizes the detection ratios of *C. parvum*, *L. pneumophila*, and MNV across different concentration ranges and provides the 50% limit of detection (i.e., LOD_50_) with McFadden’s pseudo R^2^ as estimated from the probit analysis. Note that the pseudo R^2^ for *mip* without RT-preamplification and for MNV with RT and with RT-preamplification were lower, and thus these LOD_50_ were not discussed further in this study.

**Table 1.**
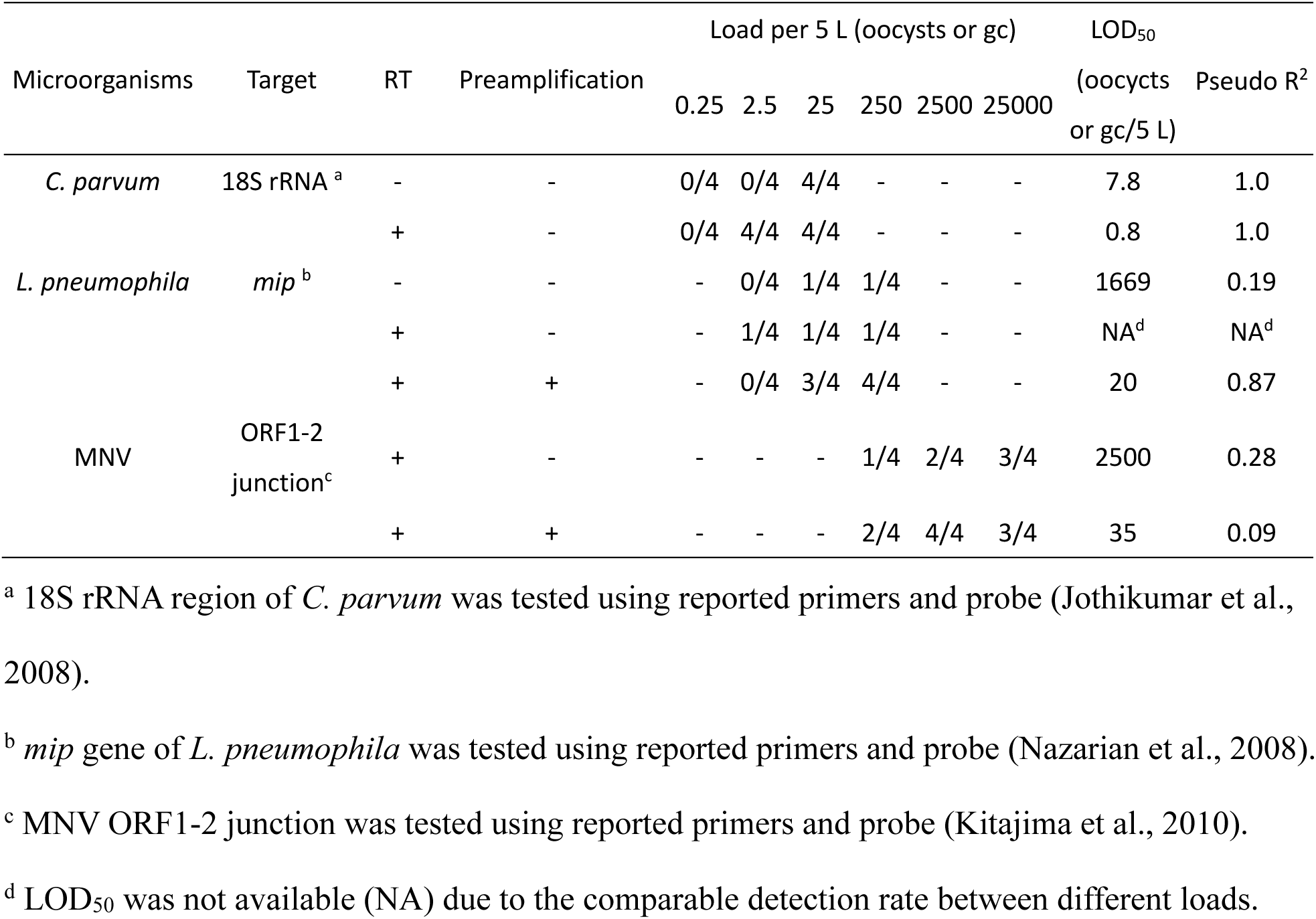
Detection ratio of *C. parvum*, *L. pneumophila*, and MNV by different ranges of concentrations and estimated LOD_50_ with McFadden’s pseudo R^2^.

| Microorganisms | Target | RT | Pre-amplification | Load per 5 L (oocysts or gc) |  |  |  |  |  | LOD<sub>50</sub><br>(oocysts or gc/5 L) | Pseudo R<sup>2</sup> |
| --- | --- | --- | --- | --- | --- | --- | --- | --- | --- | --- | --- |
|  |  |  |  | 0.25 | 2.5 | 25 | 250 | 2500 | 25000 |  |  |
| *C. parvum* | 18S rRNA <sup>a</sup> | - | - | 0/4 | 0/4 | 4/4 | - | - | - | 7.8 | 1.0 |
|  |  | + | - | 0/4 | 4/4 | 4/4 | - | - | - | 0.8 | 1.0 |
| *L. pneumophila* | *mip* <sup>b</sup> | - | - | - | 0/4 | 1/4 | 1/4 | - | - | 1669 | 0.19 |
|  |  | + | - | - | 1/4 | 1/4 | 1/4 | - | - | NA<sup>d</sup> | NA<sup>d</sup> |
|  |  | + | + | - | 0/4 | 3/4 | 4/4 | - | - | 20 | 0.87 |
| MNV | ORF1-2<br>junction<sup>c</sup> | + | - | - | - | - | 1/4 | 2/4 | 3/4 | 2500 | 0.28 |
|  |  | + | + | - | - | - | 2/4 | 4/4 | 3/4 | 35 | 0.09 |

<sup>a</sup> 18S rRNA region of *C. parvum* was tested using reported primers and probe (Jothikumar et al., 2008).
<sup>b</sup> *mip* gene of *L. pneumophila* was tested using reported primers and probe (Nazarian et al., 2008).
<sup>c</sup> MNV ORF1-2 junction was tested using reported primers and probe (Kitajima et al., 2010).
<sup>d</sup> LOD<sub>50</sub> was not available (NA) due to the comparable detection rate between different loads.

*C. parvum* was consistently detected at 25 oocysts per filter regardless of the employment of the RT. Notably, the observed concentration with RT was 662-fold higher compared with the that without RT, indicating the abundant presence of 18S rRNA compared with 18S rDNA. At lower concentrations, *C. parvum* was not detected without RT. The LOD_50_ for *C. parvum* was estimated at 7.8 oocysts/5 L without RT, whereas the employment of RT improved the LOD_50_ to 0.8 oocyst/5 L. In contrast, the detection of *mip* gene did not change substantially between with and without RT. Nevertheless, incorporating RT-preamplification markedly enhanced the sensitivity, a LOD_50_ of 20 gc/5 L. A similar improvement was observed for MNV; MNV detection rate was higher with preamplification compared with without preamplification at a load of 250 and 2,500 gc/5 L.

These results indicate that performing RT for *C. parvum* and employing RT-preamplification for the *mip* gene, as well as for MNV, resulted in the lowest LOD_50_ values, demonstrating that these approaches enhance the sensitivity.

### 3.3. Simultaneous detection of viruses, bacteria, and protozoa from river water

To test the environmental applicability of the optimized method, we applied it to the river water samples as a proof-of-concept. For each sample, one quarter of the membrane was used for detecting viable *Cryptosporidium* spp. and *Giardia* spp. by IFAT, while another quarter was used for molecular detection of these protozoa, the bacteria *L. pneumophila* and the HF183, and NoV GII.

The range of the observed concentration for each target is presented in Figure 2 and Figure 3. The temporal trends in the concentrations of each microorganism are presented in Figure S1.

**Figure 2.**
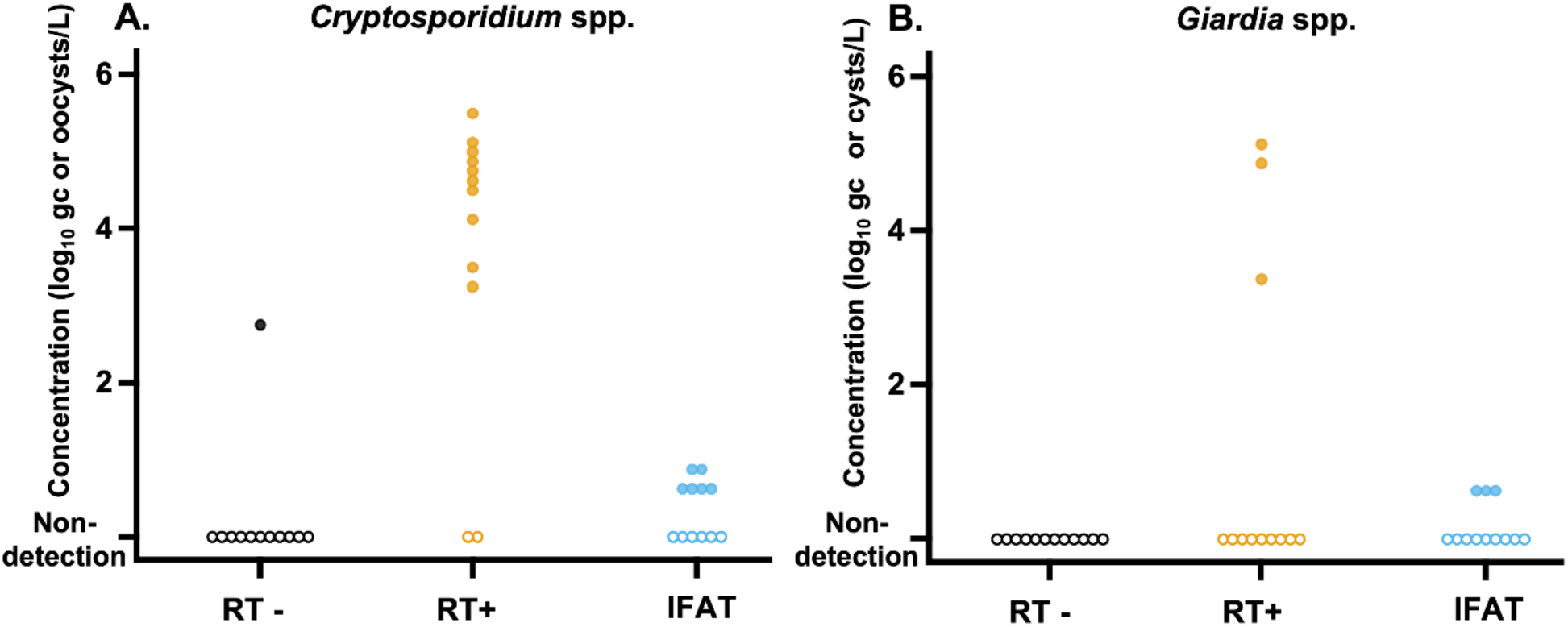
*Cryptosporidium* spp. (A) and *Giradia* spp. (B) concentrations without and with RT (RT – and RT +, respectively) and (oo)cyst concentrations measured by immunofluorescent antibody test. Empty plots indicate non-detection of the target. A total of 12 samples were tested.

**Figure 3.**
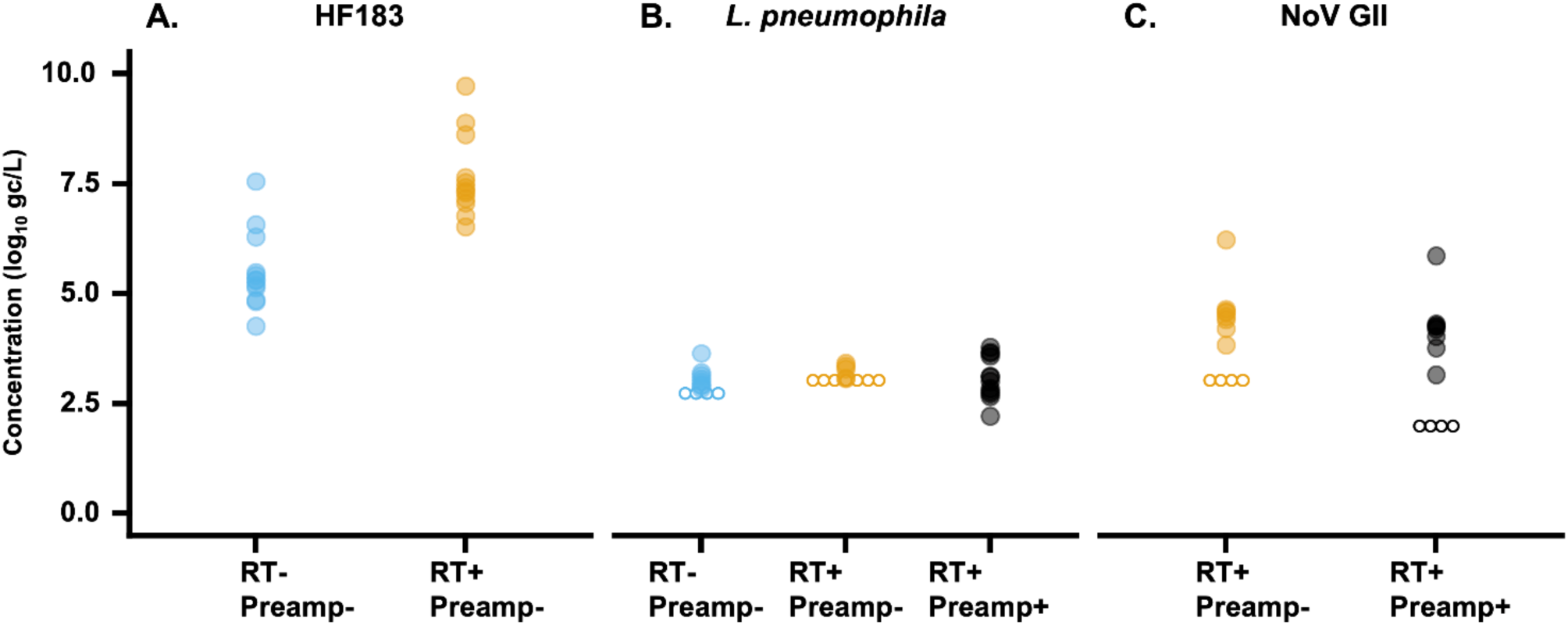
HF183 (A), *L. pneumophila mip* gene (B), and NoV GII (C) concentrations measured by different procedures, employment of RT (RT – and RT +, respectively) and preamplification (Preamp – and Preamp +, respectively). Empty plots indicate non-detection of the target. A total of 12 samples were tested.

Viable *Cryptosporidium* spp. oocysts were detected in 6 out of 12 (Figure 2A). All the detections were close to the theoretical limit of detection, single (oo)cysts per 250 mL. Notably, *Cryptosporidium* spp. was detected in 10 of 12 samples with RT, compared with only 1 of 12 samples without RT (Fisher’s exact test, *P <* 0.01). In addition, the observed concentrations for *Cryptosporidium* spp. with RT ranged from 3.2 to 5.5 log gc/L, and are higher than those without RT by 0.4 to 2.8 log gc/L. Similarly, while *Giardia* spp. were detected in 3 of 12 samples with RT, *Giardia* spp. was not detected in all samples without RT (Figure 2B). The similar or even higher detectability for protozoan rRNA compared with IFAT is likely due to a limitation of PCR, which detects both viable and non-viable organisms, whereas IFAT selectively detects viable (oo)cysts. These results indicate that targeting 18S rRNA by performing RT enhances the detectability of protozoa.

HF183 was detected from all samples regardless of the employment of RT (Figure 3A). The concentration without RT, ranged from 4.3 to 7.6 log gc/L (Figure 3A and Figure S1C). Similar to the case of protozoan 18S rRNA, the concentrations of HF183 markers with RT are higher in samples with 2.2 log higher than HF183 without RT on average, suggesting that targeting the rRNA still holds a potential to improve the sensitive detection of HF183, especially in settings with low microbial contamination level, such as drinking water sources.

In contrast, the detection of *L. pneumophila mip* gene did not substantially differ between the two procedures, RT− Preamp− (8/12) and RT+ Preamp− (5/12) (Figure 3B). Inclusion of RT-preamplification allowed for the detection of *mip* gene marker under the LOD of RT-qPCR (see the data points in Oct 2023 and Jan 2024 in Figure S1), confirming the utility of RT-preamplification to enhance sensitivity as discussed in the section before. Importantly, the use of RT-preamplification did not alter the observed concentrations, indicating that this step does not interfere with the measurement.

NoV GII was detected from 8 out of 12 samples, irrespective of whether preamplification was applied (Figure 3C). This comparable detection rate is possibly because the NoV GII concentration for the four points is below the LOD for RT+ Preamp+ (Figure S1E), and thus the effect of RT-preamplification was not pronounced. Aligning with the data on *L. pneumophila*, the observed NoV GII concentration did not differ between with and without RT-preamplification.

Overall, these results demonstrate that targeting rRNA for protozoa and bacteria markedly improves detection sensitivity. Moreover, for targets where only a single or a few copies are present per organism, such as bacterial genomic DNA or viral genomic RNA, RT-preamplification serves as a valuable tool to further enhance sensitivity without introducing quantification bias.

### 3.4. Species identification of *Giradia* spp

To identify the species of *Giardia* spp., three real-time PCR products that tested positive were subjected to nanopore sequencing. Each sample yielded a contig that perfectly matched the 18S rRNA gene of *Giardia duodenalis* (accession no. AF199446), indicating that the detected *Giardia* included the human-infectious *G. duodenalis*. These results highlight the advantage of molecular methods in providing more detailed species-level characterization compared to conventional IFAT.

## 4. Discussion

This study showed that the membrane-adsorption and direct nucleic acid extraction allow for simultaneous and sensitive detection of protozoa, bacteria, and viruses from a single membrane filter used to process environmental water samples.

Firstly, our optimization experiment revealed that the use of a mixed cellulose ester membrane with a pore size of 0.8 μm, along with supplementation of magnesium chloride in water, recovered protozoa, bacteria, and viruses most efficiently. In particular, the supplementation with magnesium chloride improved virus recovery by up to 120-fold for the mixed cellulose ester membrane. This improvement likely occurs because magnesium chloride promotes the adsorption of viruses on the mixed cellulose ester filter, whose pore size is larger than viruses, and is consistent with a previous study (Lukasik et al., 2000). Although the nylon membrane recovered viruses to some extent even without ion supplementation and was consistent with a previous report (Oshima et al., 1994), their lower water permeability (data not shown) makes them less suitable for processing large volumes of water. The effect of pore size on the recovery for mixed cellulose ester membranes was not pronounced within a range of pore sizes used in this study, with a 1.5-fold difference for *C. parvum* and *L. pneumophila* and with a 2.1-fold difference for MNV. This suggests that the use of membrane filter with a larger pore size could be an option when the water samples are prone to causing membrane clogging. The optimized method is analogous to the EPISENS-M developed for detecting SARS-CoV-2 (Ando et al., 2023; Nakaso et al., 2024) under a context of wastewater-based epidemiology. This suggests that EPISENS-M can be used as a general procedure to detect pathogens regardless of the type of water samples for water utilities.

Moreover, the method achieved >73% recovery of *C. parvum*, *L. pneumophila*, and MNV and allows for highly sensitive detection. Notably, the sensitivity for *C. parvum* was equivalent to conventional microscopy-based IFAT. A key factor in achieving this sensitivity was performing RT to target not only 18S rDNA but also 18S rRNA molecules. Since a single *C. parvum* oocyst contains four sporozoites, each of which has two to five rDNA (Huang et al., 2024), the total number of rDNA per oocyst is limited to only eight to twenty. In contrast, rRNA was found to be present in quantities up to hundreds of times greater than rDNA as detailed in section 3.2. Given that qPCR analyzes only a subsample of nucleic acid extracts, it is plausible that targeting the more abundant rRNA enhances sensitivity, which is consistent with a previous work reporting an advantage of the employment of RT for protozoa detection (Kishida et al., 2013). A similar benefit was observed for HF183, suggesting that rRNA-based detection may offer a broadly applicable strategy for the sensitive detection of both protozoa and bacteria. It should be cautious that rRNA expression is expected to vary largely between different oocysts and cells. In fact, the rRNA concentration of *Cryptosporidium* spp. and *Giardia* spp. differ by up to 2-log across different samples even though the (oo)cyst concentration was at a comparable level, ranging from zero to two (oo)cyst per 250 mL, for all the samples as shown in Figure 2. Therefore, the decision to employ RT should be based on the intended application: RT is recommended when sensitive detection is needed for monitoring water quality deterioration, yet not recommended when accurate pathogen quantification is required for quantitative microbial risk assessment (Pecson et al., 2022).

It is noteworthy that this study also preliminarily tested the applicability of 16S rRNA-based detection of *L. pneumophila* using a developed assay (Donohue et al., 2014), but was excluded from the analysis in this study due to the non-specific reaction occurring in NTC that underwent RT or RT-preamplification. Contrary to the improvements observed with rRNA targeting, focusing on the mRNA of *mip* gene of *L. pneumophila* did not significantly enhance sensitivity. Given that *mip* gene is involved in resistance to intracellular killing and multiplication (Cianciotto and Fields, 1992), it would not be surprising if *mip* mRNA is merely expressed in aquatic environments, such that performing RT did not have large effect on the sensitivity.

An alternative strategy to improve sensitivity is the application of RT-preamplification. In the settings of the present study, the ESV for qPCR was 1.25 or 2.5 μL of nucleic acid extract for RNA or DNA target, respectively, while that for RT-preamplification was 13.5 μL. This increase in ESV improved the sensitivity of detection, particularly for low-copy targets such as viral RNA and the single-copy *mip* gene (Table 1 and Figure 3B). Importantly, the observed measurements obtained by RT-qPCR and RT-preamplification-qPCR were comparable (Figure 3B and C), suggesting that the RT-preamplification does not interfere with the ability for quantification.

Current regulatory guidelines typically require that pathogens be confirmed as “negative” or below a certain threshold rather than precise quantification (e.g., negative for *E. coli* in 100 mL of test water). Given the sensitivity achieved in this study, our method may serve as a robust ‘screening’ tool for confirming the absence or acceptable low levels of protozoa, bacteria, and viruses in environmental samples. However, it is important to note that molecular methods cannot distinguish between viable and non-viable organisms. Therefore, when positive results are obtained, confirmatory testing for viability or infectivity remains essential. Notably, the concentration step in our method is analogous to those employed in conventional IFAT and culture-based virus detection protocols (Haramoto et al., 2012; Katayama et al., 2002). Moreover, the remaining portions of the membrane filter can be directly subjected to these traditional methods, as illustrated in the graphical abstract. This methodological compatibility offers reassurance to water utilities that may be hesitant to transitioning from culture- or IFAT-based assays to the molecular-based workflow.

## 5. Conclusion

This study demonstrates that membrane adsorption with direct nucleic acid extraction is a sensitive approach for simultaneously detecting protozoa, bacteria, and viruses in environmental water samples. Optimizing the process by using a mixed cellulose ester membrane (0.8 μm) and magnesium chloride supplementation significantly enhanced pathogen recovery for the three pathogens. Employment of RT to target not only rDNA but rRNA significantly enhanced sensitivity for protozoa and bacterial indicators, while RT-preamplification improved detection limits for targets present in low copy numbers, such as viral and single copy-bacterial genomes. Environmental testing using river water samples further validated the approach, showing that the sensitivity of molecular detection can be comparable to conventional IFAT for protozoa. Overall, this method enables comprehensive ‘screening’ for protozoa, bacteria, and viruses in environmental waters. It reduces the need for labor-intensive techniques like IFAT and culture-based virus detection, thereby lowering the workload for water utilities during pathogen monitoring.

## 6. Acknowledgment

This study was supported by the JSPS through Grant-in-Aid for Scientific Research (A) (grant number 24H00326 and 25H00755) and for Young Scientists (grant number 24K17379), AMED under Grant Number JP223fa627001, the River Fund of The River Foundation, Japan under Grant Number 2023-5311-005, Kurita Water and Environmental Foundation (grant number 23H036, 24K023, and 25K023), Next Generation Water Supply Research Incentive Program for Young Researchers supported by Kubota, and the Environment Research and Technology Development Fund (grant number JPMEERF20255RB1) of the Environmental Restoration and Conservation Agency provided by Ministry of the Environment of Japan. During the preparation of this work, the authors used ChatGPT 4o mini solely for improving readability and language in accordance with the journal’s policies. After employing this tool, the author reviewed and edited the content as needed and takes full responsibility for the content of the publication.

## 7. Data Availability

All data discussed in this manuscript will be accessible on zenodo upon the acceptance of the manuscript.

## 8. CRediT Authorship Contribution Statement

**Shotaro Torii:** Formal analysis, Conceptualization, Investigation, Methodology, Validation, Writing – Original draft, **Masaaki Kitajima:** Methodology, Writing – review and editing, **Eiji Haramoto:** Methodology, Writing – review and editing, **Ryota Gomi:** Methodology, Writing – review and editing **Kumiko Oguma:** Methodology, Writing – review and editing, **Hiroyuki Katayama**: Writing – review and editing

## S1: Digital droplet PCR for the enumeration of the gBlocks

The gBlocks Gene Fragments purchased were enumerated by digital droplet PCR (ddPCR). The gBlocks were resuspended with Tris-EDTA buffer to bring the final concentration, reported from the manufacturer, to 10^10^ gc/μL. After being diluted by 5×10^6^-fold, the gBlocks are used as a template for ddPCR. ddPCR was performed in a total reaction volume of 20 μL, which consisted of 2 μL of the template and 18 μL of the reaction mixture. The reaction mixture consisted of 10 μL of ddPCR™ Supermix for Probes, 6 μL of nuclease free water, and 18 pmol each of forward and reverse primers, and 9 pmol of probe. The same primers and probe as real-time PCR were used. Droplets were generated using the QX200 Droplet Generator (Bio-Rad) and amplified using the C1000 Touch Thermal Cycler (Bio-Rad). After amplification, droplets were analyzed using the QX200 droplet reader (Bio-Rad) and the Quantasoft Analysis Pro Software. Table S1 shows the observed concentration of the gBlocks. gBlocks were diluted according to the observed concentration by ddPCR and stored at −80℃ until qPCR.

## S2: Nanopore sequencing of real-time PCR products of *Giardia* spp

For species identification of *Giardia* spp., real-time PCR products were sequenced. Gel electrophoresis revealed multiple bands of similar amplicon lengths, prompting us to choose nanopore sequencing for downstream analysis instead of direct sequencing, which was used in the original study (Jothikumar et al., 2021). Briefly, three real-time PCR products that tested positive with the *Giardia* spp. assay were purified and submitted for nanopore sequencing using the Premium PCR sequencing service from Plasmidsaurus (USA). This service employs the ligation sequencing Kit v14 (Oxford Nanopore Technologies) and R10.4.1 flow cells (Oxford Nanopore Technologies), enabling assembly accuracy of Q50 at a sequencing depth of 10–20× according to the manufacturer. A total of 5,000 reads were obtained for each sample. They were trimmed of primer sequences of *Giardia* spp., and de novo assembled. The top five high-coverage contigs, each of which assembled a total of 120−790 reads, were blasted using Geneious Prime® 2025.1.2.

---

**Figure S1.**
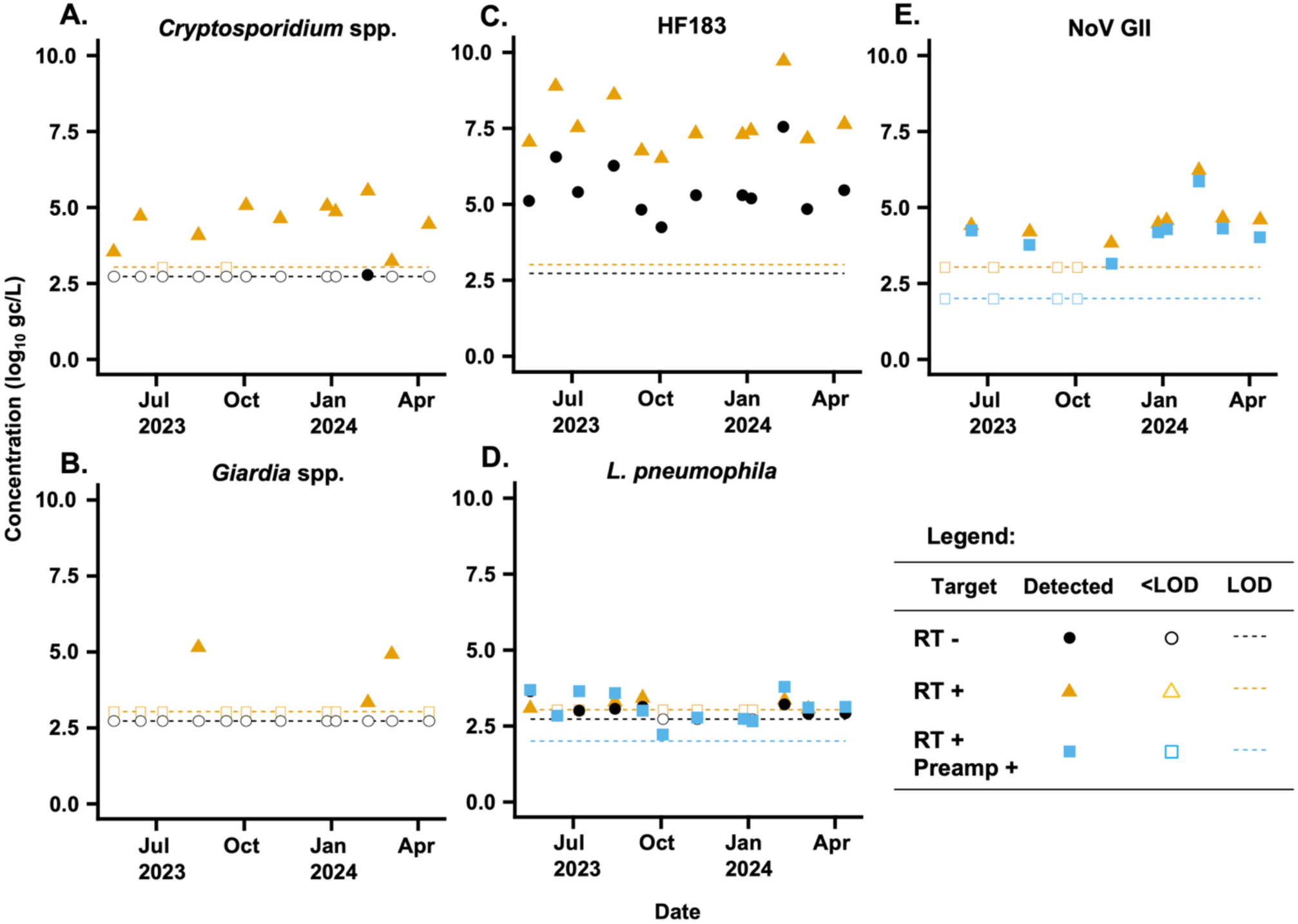
Concentration of *Cryptosporidium* spp. (A), *Giardia* spp. (B), HF183 (C), *L. pneumophila* (D), and NoV GII (E) measured by real-time PCR in the Tama River. Black circular symbols indicate concentrations obtained without the RT process (i.e., RT –), orange triangles represent concentrations with the RT process (i.e., RT +), and blue squares denote concentrations after one-step RT-preamplification (i.e., RT + Preamp +). Open symbols represent results below the detection limits, and dashed lines indicate the LOD for each procedure. The LOD was three gene copy number per reaction; 3 gc per 2.5 μL of nucleic acid extract RT –, 3 gc per 1.25 μL of nucleic acid extract for RT +, 3 gc per 13.5 μL of nucleic acid extract for RT+ Preamp+.

**Table S1.**
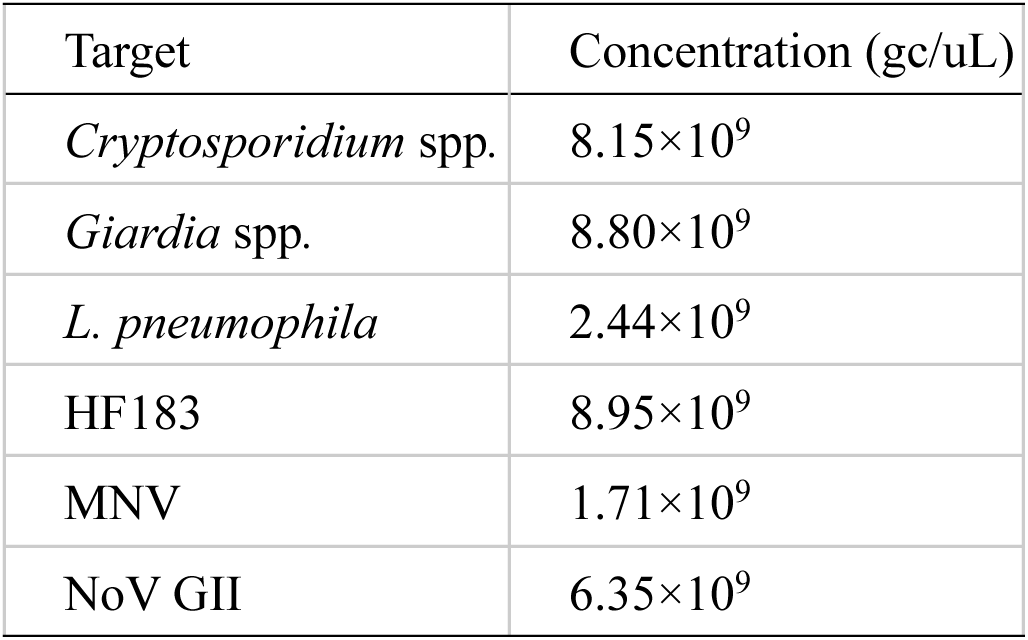
Observed concentration of 10^10^ gc/μL of gBlocks by ddPCR.

**Table S2.**
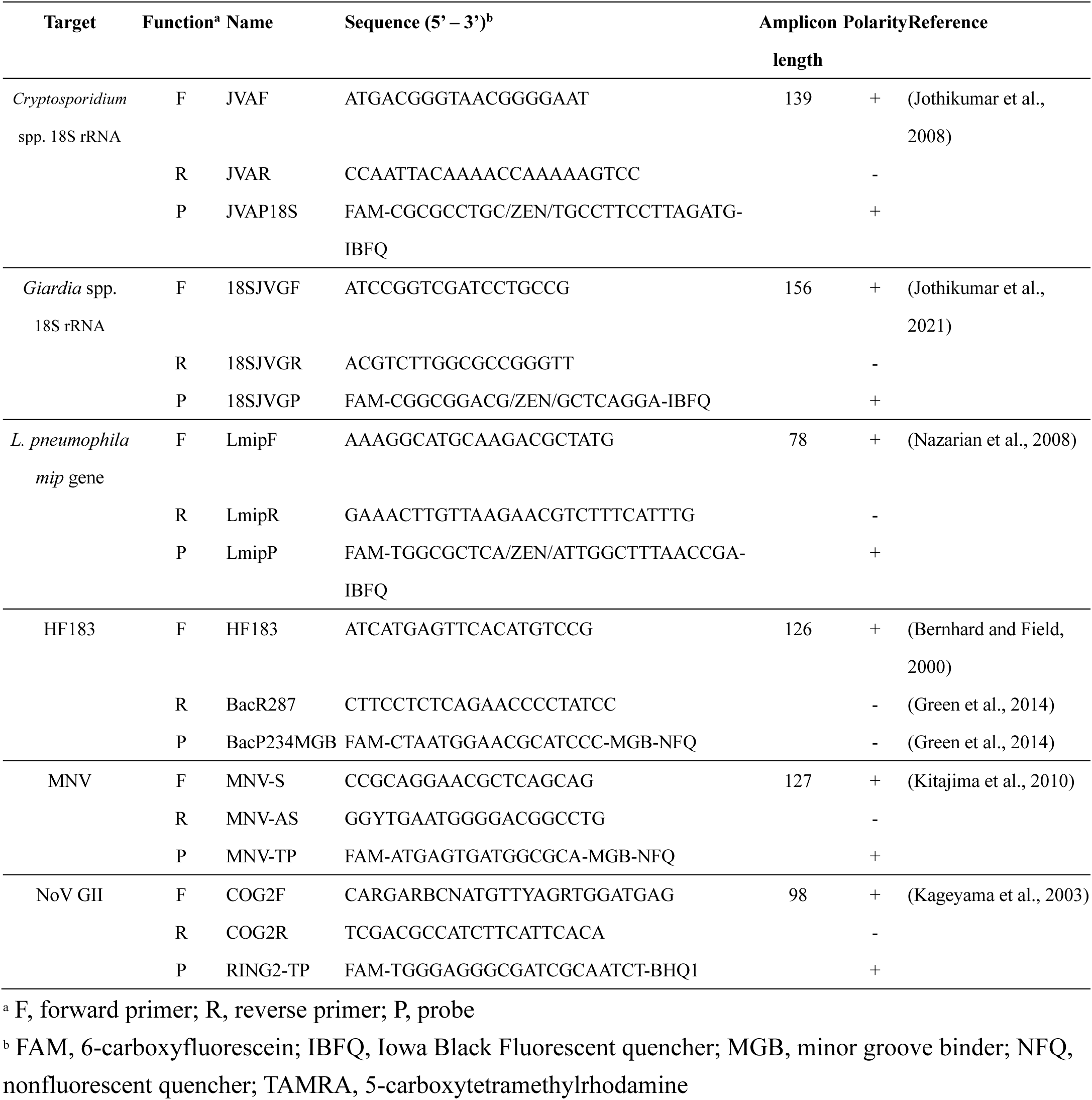
Primers and probes used for real-time PCR.

