## Supplementary material for "Simultaneous detection of protozoa, bacteria, and viruses from environmental water through membrane-adsorption followed by direct nucleic acid extraction": SI

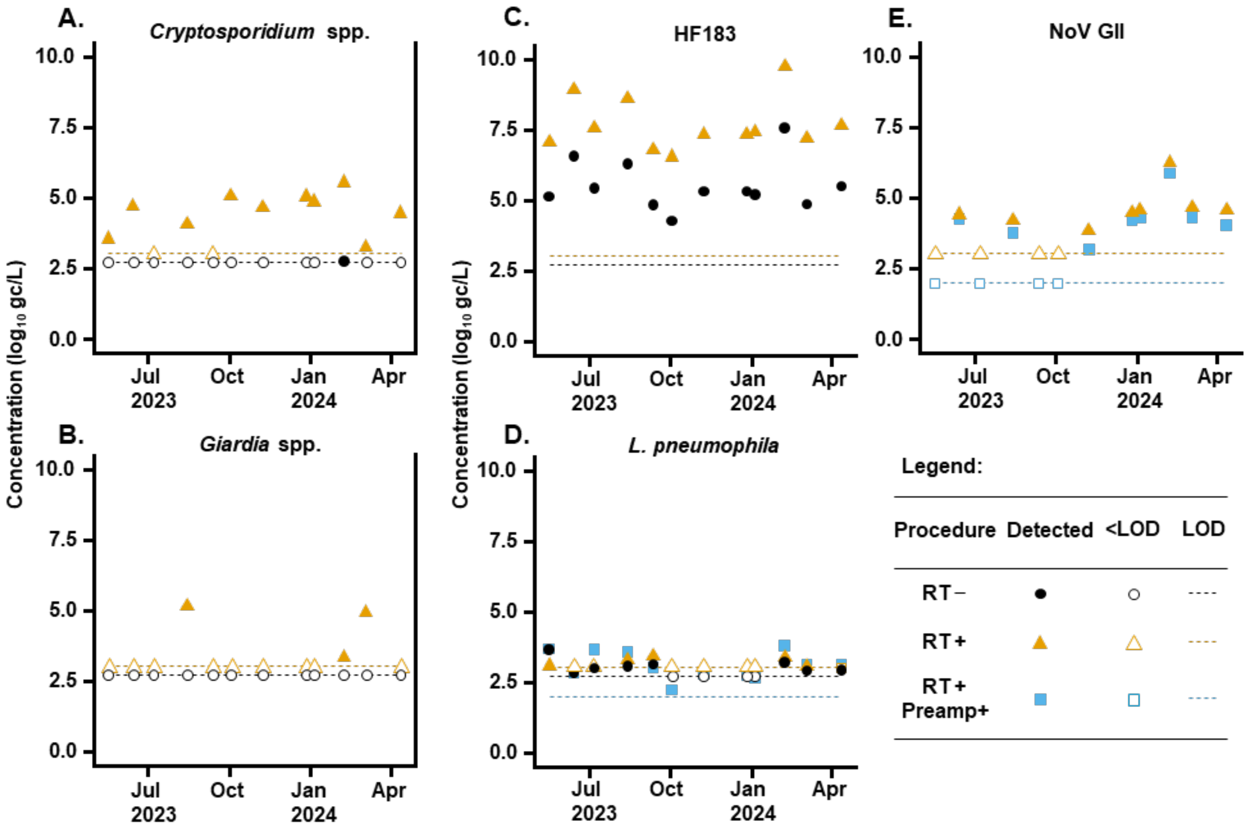


Figure S1 Concentration of *Cryptosporidium* spp. (A), *Giardia* spp. (B), HF183 (C), *L. pneumophila* (D), and NoV GII (E) measured by real-time PCR in the Tama River. Black circular symbols indicate concentrations obtained without the RT process (i.e., RT –), orange triangles represent concentrations with the RT process (i.e., RT +), and blue squares denote concentrations after one-step RT-preamplification (i.e., RT + Preamp +). Open symbols represent results below the detection limits, and dashed lines indicate the LOD for each procedure. The LOD was three gene copy number per reaction; 3 gc per 2.5 µL of nucleic acid extract RT –, 3 gc per 1.25 µL of nucleic acid extract for RT +, 3 gc per 13.5 µL of nucleic acid extract for RT+ Preamp+.

Table S1 Observed concentration of $10^{10}$ gc/µL of gBlocks by ddPCR.

| Target | Concentration (gc/uL) |
|---|---|
| *Cryptosporidium* spp. | $8.15 \times 10^9$ |
| *Giardia* spp. | $8.80 \times 10^9$ |
| *L. pneumophila* | $2.44 \times 10^9$ |
| HF183 | $8.95 \times 10^9$ |
| MNV | $1.71 \times 10^9$ |
| NoV GII | $6.35 \times 10^9$ |

Table S2 Primers and probes used for real-time PCR

| Target | Function[a] | Name | Sequence (5' – 3')[b] | Amplicon length | Polarity | Reference |
|---|---|---|---|---|---|---|
| *Cryptosporidium* spp. 18S rRNA | F | JVAF | ATGACGGGTAACGGGGAAT | 139 | + | (Jothikumar et al., 2008) |
| | R | JVAR | CCAATTACAAAACCAAAAAGTCC | | - | |
| | P | JVAP18S | FAM-CGCGCCTGC/ZEN/TGCCTTCCTTAGATG-IBFQ | | + | |
| *Giardia* spp. 18S rRNA | F | 18SJVGF | ATCCGGTCGATCCTGCCG | 156 | + | (Jothikumar et al., 2021) |
| | R | 18SJVGR | ACGTCTTGGCGCCGGGTT | | - | |
| | P | 18SJVGP | FAM-CGGCGGACG/ZEN/GCTCAGGA-IBFQ | | + | |
| *L. pneumophila mip* gene | F | LmipF | AAAGGCATGCAAGACGCTATG | 78 | + | (Nazarian et al., 2008) |
| | R | LmipR | GAAACTTGTTAAGAACGTCTTTCATTTG | | - | |
| | P | LmipP | FAM-TGGCGCTCA/ZEN/ATTGGCTTTAACCGA-IBFQ | | + | |
| HF183 | F | HF183 | ATCATGAGTTCACATGTCCG | 126 | + | (Bernhard and Field, 2000) |
| | R | BacR287 | CTTCCTCTCAGAACCCCTATCC | | - | (Green et al., 2014) |
| | P | BacP234MGB | FAM-CTAATGGAACGCATCCC-MGB-NFQ | | - | (Green et al., 2014) |
| MNV | F | MNV-S | CCGCAGGAACGCTCAGCAG | 127 | + | (Kitajima et al., 2010) |
| | R | MNV-AS | GGYTGAATGGGGACGGCCTG | | - | |
| | P | MNV-TP | FAM-ATGAGTGATGGCGCA-MGB-NFQ | | + | |
| NoV GII | F | COG2F | CARGARBCNATGTTYAGRTGGATGAG | 98 | + | (Kageyama et al., 2003) |
| | R | COG2R | TCGACGCCATCTTCATTCACA | | - | |
| | P | RING2-TP | FAM-TGGGAGGGCGATCGCAATCT-BHQ1 | | + | |

[a] F, forward primer; R, reverse primer; P, probe

[b] FAM, 6-carboxyfluorescein; IBFQ, Iowa Black Fluorescent quencher; MGB, minor groove binder; NFQ, nonfluorescent quencher; TAMRA, 5-carboxytetramethylrhodamine
